# The microbiome of an amphibious plant shifts dramatically across the host life cycle

**DOI:** 10.1101/2023.08.20.552558

**Authors:** Jorge A Mandussí Montiel-Molina, Jason P Sexton, J Michael Beman, A Carolin Frank

## Abstract

Interactions between plants and their associated microbiomes are thought to enhance the capacity of the host plant to overcome extreme conditions, yet the significance of microbially mediated stress tolerance in most plant species and ecosystems remains unknown. For the first time, we examined the microbiome of the amphibious plant *Eryngium castrense* (Apiaceae) inhabiting Mediterranean climate ephemeral wetlands in California. *Eryngium castrense* has the capacity to survive as both an aquatic and terrestrial plant, thereby living under contrasting extremes of water stress. Whether plant-associated microbial communities are also affected by such changes, and what ecological role they play while inhabiting amphibious plants, is an underexplored topic of plant-microbial interactions in natural and artificial systems. We amplified and sequenced 16S rRNA genes from bacteria and archaea to examine microbial communities associated with roots and shoots over the plants’ full life cycle. We observed that the microbiome changes from the aquatic stage to the terrestrial stage, and that roots and shoots represent distinct habitats within the plant host ecosystem. When compared with soil and water column samples, plant samples retained a unique, differentiated core microbiome. Taxa located in the roots during the terrestrial stage were linked with potential functions such as nitrogen acquisition, sulfur assimilation, and resistance to heavy metals, whereas aquatic roots held potential phythoparasites. Overall, our results provide new insights into symbiotic relationships in plants subject to stress-related to water saturation and deficiency.

## 1 Introduction

Symbiosis with microbes can improve plant health and habitat adaptation in natural and artificial environments, with several plant traits co-regulated by the associated microbiome and the plant’s genome (Rodriguez, et al 2009; Morelli, et al 2020). Research focusing on symbiotic associations can also enhance ecological theory and may even lead to the development of new biotechnology (Redman et al, 2022). Among symbionts, microbial inhabitants of the rhizosphere and phyllosphere are considered epiphytes, whereas microbes residing within plant tissues (the endosphere) are considered endophytes. These different types of symbionts can also occupy different within-plant habitats (compartments)—such as roots, shoots, and leaves—establishing beneficial associations with their host plants (Carrell & Frank, 2014; Fan, et al 2020). Selection imposed by plant compartments strongly shapes the diversity and composition of the microbiota. Different plant tissues such as leaves, roots, or flowers can harbor unique taxa, modulating microbe-microbe interactions, due to both their structural differences and exposure to soil versus air and feeding back to host fitness (Fitzpatrick et al, 2020). Nevertheless, sometimes bacterial communities in leaves and roots can be surprisingly similar (Bai et al, 2015; Van der Heijden et al, 2016). While plant microbiomes are usually dominated by Proteobacteria (particularly those of the α and β classes), and other major groups include Actinobacteria, Firmicutes, Bacteroidetes, Planctomycetes, Verrucomicrobia, and Acidobacteria (Turner et al, 2013), they can be variable and complex. In addition, existing evidence suggests that variation in the abiotic and biotic environment can also exert direct and indirect effects on plant-associated microbiota; climate-driven geographical variation can drive the composition of a microbiome (Oono, et al 2017; Harrison & Griffin, 2019). The plant microbiome therefore holds interesting parallels to other environmental systems and their microbiomes—for instance, the gut microbiome or the Earth microbiome—with conceptually similar questions regarding horizontal transfer of symbionts, diversity drivers, and influence in host niche expansion and plasticity (Redman et al, 2011; Redman et al, 2022).This is of importance in a global change context, for example, as global temperature increases and variation in both abiotic and biotic factors can indirectly shape plant microbiota through plant responses (Suryanarayanan & Shaanker, 2021; Ware, et al 2021). Given the known importance of plant microbiomes and their variations, examining the communities present on and within different and unique plant species has the potential to provide new insights.

Vernal pools—ephemeral bodies of water—provide habitat for specialized plants and animals, and represent an extreme version of the climatic variation characteristic of many other ecosystems. With seasonal water saturation during winter and gradual desiccation conditions the rest of the year, vernal pools (en español: *charcas vernales*) trigger evolutionary processes of dormancy and amphibious life cycles with unique morphological traits. Many plant lineages have independently adapted to vernal pools conditions, including aquatic plants that have adapted to desiccation conditions (e.g. *Isoetes howellii*), and plants whose ancestors were terrestrial taxa that became aquatic (e.g. *Eryngium spp*.). Typically the aquatic morphology, ‘grass-like tufts’, is emblematic of the aquatic period of the pools in winter and early springtime. Later in the season, after evaporation and total desiccation of pools, plants acquire a terrestrial morphology that is spiny and resembles more woody herbaceous plants (Keeley, 1999). However, little is known about the microbiomes of these fascinating plants.

Interactions between plants and their associated microbes can provide higher fitness under extreme environmental conditions (Baedke, et al 2020; Trivedi et al, 2020) and in the context of future of agriculture practices (i.e. global change, artificial life support systems), achieving the ability to assist plants in rapidly adapting or acclimating signifies a milestone in the development of resilient crops. In this vein, amphibious plants, inhabiting ephemeral wetlands such as vernal pools are ideal study cases, as these plants tolerate wide extremes of water availability and other abiotic stress. These plants and their associated microbial symbionts can provide key insights into stress tolerance. To our knowledge, this study is the first of its kind in exploring amphibious plants and their associated microbiome. We examined *Eryngium castrense* (Apiaceae), colloqially known as the Great Valley button celery (Figure 1). This plant species is endemic to the California Central Valley, with several amphibious members of this genus in other regions of California, USA, and Baja California, Mexico in other vernal pool complexes.

**Figure 1.**
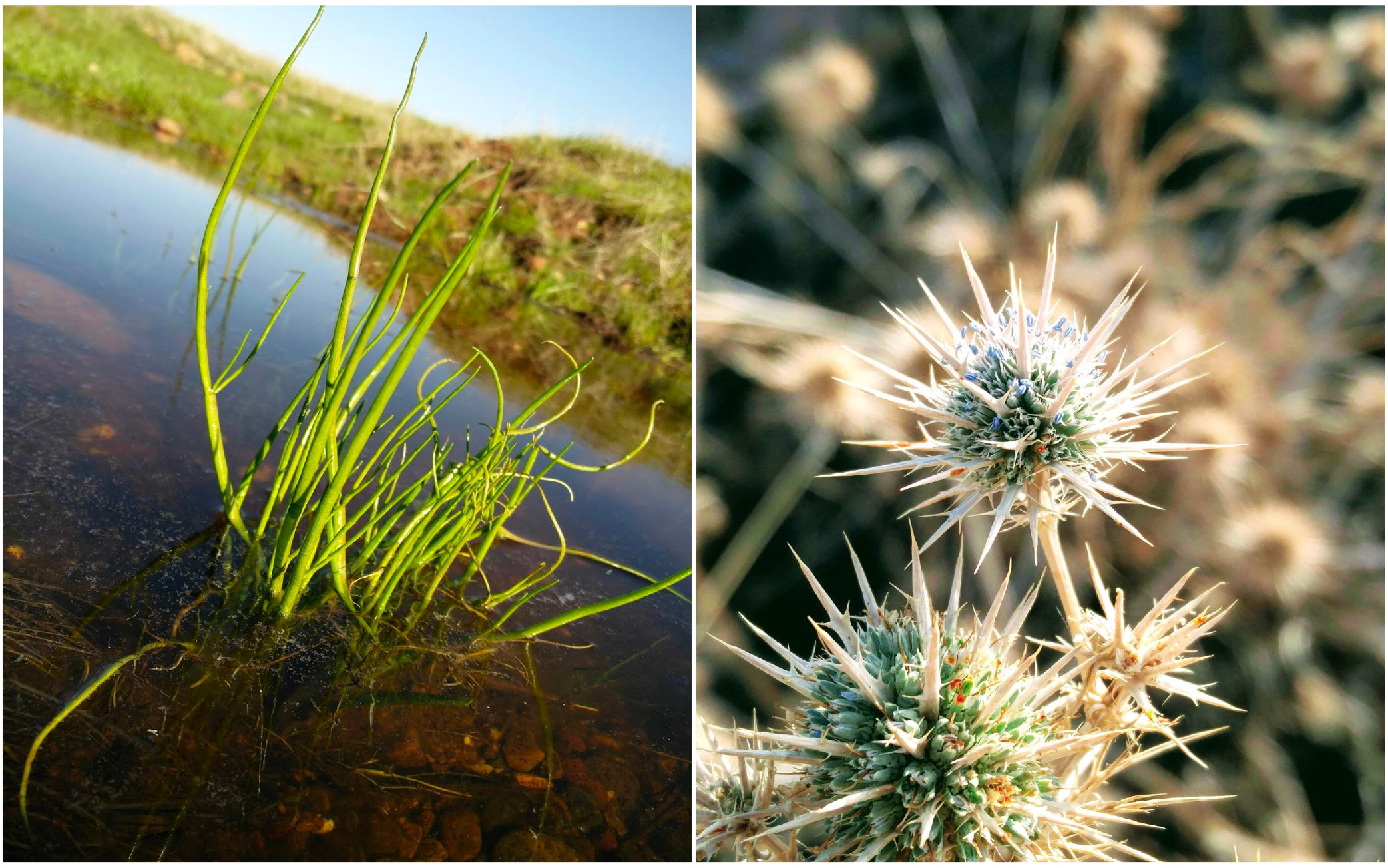
*Eryngium castrense* Jeps. is a California native plant, specialized to live under aquatic and desiccated environmental conditions. It grows in California vernal pools, which are temporary, Mediterranean-climate wetlands **(A)** “lsoetoide” aquatic morphology; **(B)** terrestrial morphology - “Spiny” weed.

In this present study, we investigated the dynamics of the microbiome (specifically bacterial endophytes) within the amphibious vernal pool plant *E. castrense*. Our first objective was to compare the differences in endophytic community diversity and composition between the aquatic and terrestrial stages of these plants. We hypothesized that if the microbial community plays a role in plant stress tolerance during extreme stages, we would observe shifts in the community composition and diversity aligned with aquatic and desiccation stages. Additionally, we examined the differentiation within the host habitat by comparing the root and shoot compartments. We aimed to determine whether these plant compartments harbor distinct microbial communities, reflecting the different above-ground and below-ground needs of the plant. We also examined community assembly at the ecosystem level, focusing on differences in composition and diversity between endophytic communities and immediate plant surroundings, including the soil surrounding the roots and the water column that is in contact with the shoots. Our objective was to identify a unique core microbiome in the soil, water, and plants, indicating the presence of specific ecological niches.

Finally, we investigated the influence of the surrounding environment as a filtering agent for community composition. We aimed to understand the extent to which the microbiome of amphibious plants is shaped by the plant host, environmental selection, or dispersal, considering the influence of landscape heterogeneity. Our overarching hypothesis motivating this work was that the microbial community is relevant to plant stress tolerance across the extreme stages in vernal pools, with microbes supporting critical functions when the plant host is subject to desiccation and water saturation stresses. We anticipated that compartmentalization within the plant, based on tissue differentiation, would result in niche specificity, with distinct microbial functions within shoots and roots. Additionally, we expected that landscape properties would influence the composition of the plant microbiome.

## 2 Methods

### 1.1 Study area

The study area is located in the heart of the San Joaquin Valley, California, USA, a 400 kilometers long and 80 kilometers wide open are, flanked on the east by the Sierra Nevada and on the west by the Coast Range mountains. The climate of the region is Mediterranean, characterized by cool, wet winters and hot, dry summers; annual rainfall generally ranges from 230-380 mm with 90% of the precipitation occurring from November to April. The western and eastern boundaries of the region delimit a distinct topographic and biogeographic unit of undulating terrain, from above the historic San Joaquin River floodplain, to the base of the Sierra Nevada foothills. This area supports the largest block of unfragmented vernal pool habitat remaining in California, characterized by low-slope basins with undulating *mima mound* topography that typically supports a high density of vernal pools (Vollmar, 2002). The sampling was performed at the University of California, Merced Vernal Pools and Grassland Reserve (MVPGR) (Figure 2), adjacent to the main campus of the University of California, Merced. The MVPGR is tightly managed for conservation. However, we hypothesized that development effects may be stronger on the west side of the reserve facing campus and the city of Merced. On the western boundary of the reserve, activities derived from land management, such as high cattle concentration, disturbances derived from urbanization, such as concrete dust, light pollution, noise or, activities derived from the UC campus such as visitors, trash, and motor vehicles, may lead to alterations of the natural landscape and create an anthropogenic impact zone.

**Figure 2.**
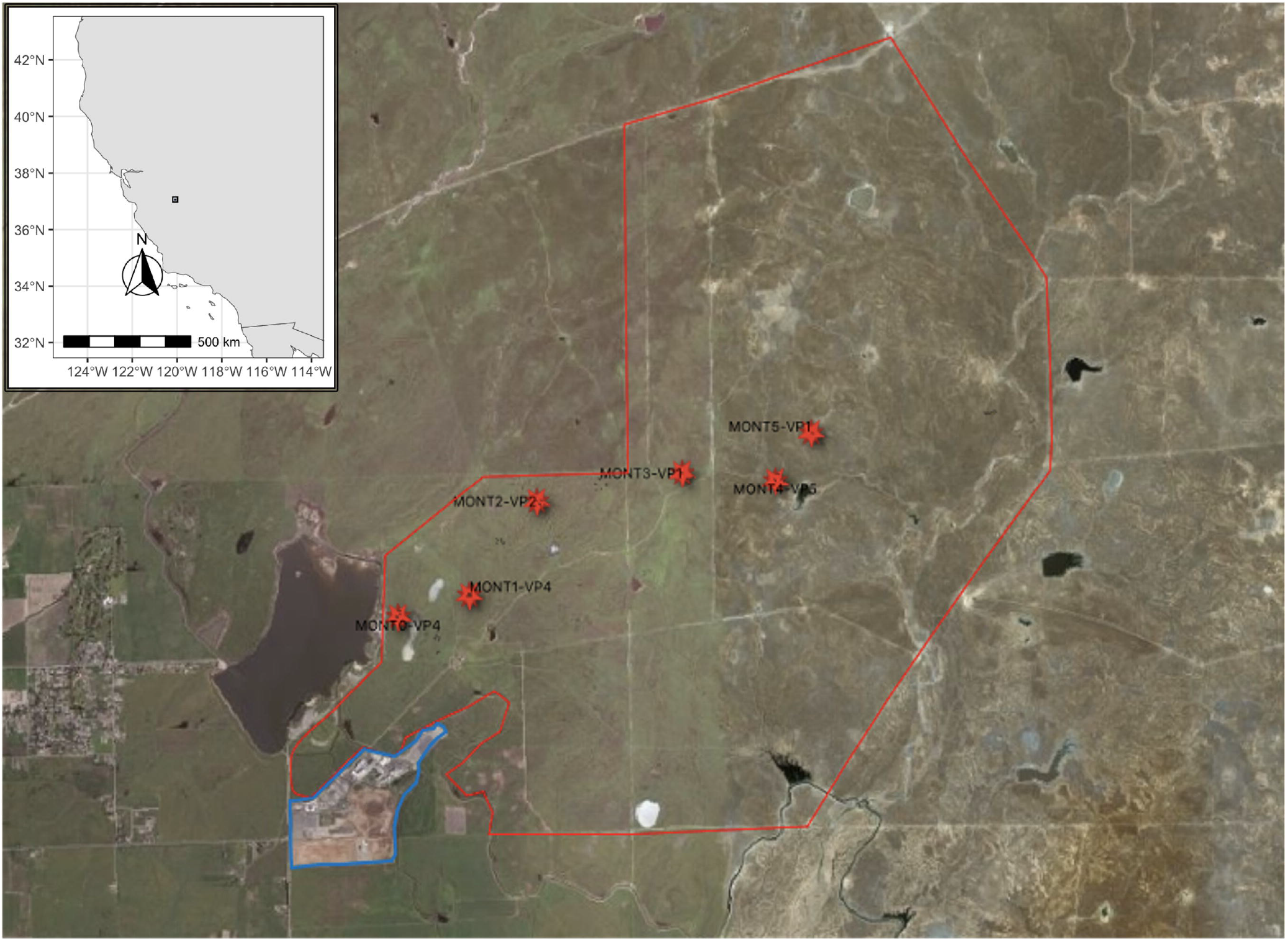
Map of the study site. UC-Merced Vernal Pools and Grassland Reserve (MVPGR) is delimited by the red polygon. The reserve is adjacent to the University of California campus, delimited with a blue polygon. Red marks indicate the location of vernal pools monitored for the studied microbes.

### 1.2 Sampling

During the first months of life development, *Eryngium castrense* resembles aquatic grasses (“isoetoide” morphology, Fig 1a), extended above the vernal pool water surface. After the pool dries out, it becomes weedy, with sharp, spiny leaves (Fig 1b), flowering throughout the hot summer (Preston, et al 2023). This plant species has well-differentiated roots and shoots, and a differentiated phenotype as a result of the metamorphosis from aquatic to terrestrial morphology, corresponding to soil flooding and desiccation. *E. castrense* is a glabrous, ascending herbaceous plant, multi-branched and spiny, with lanceolate, deeply pinnate leaves (Baldwin & Goldman, 2012; Preston, et al 2023). This plant species is abundant inside the limits of UC Merced’s Vernal pools and grassland reserve and many vernal pools in California’s Central Valley.

Collections of specimens of *E. castrense* were made at five vernal pools (Table 1). Sites were selected by following a five-kilometer transect, starting from the western boundary of the reserve, adjacent to the UC Merced campus, moving east, each site one kilometer apart. Collections were made in two seasons; during the aquatic period, which occurred in winter 2018 and comprised 20 plant specimens (four individuals per pool); followed by a collection of 25 specimens (five individuals per pool) in late spring 2018, for a total of 45 plant specimens. This sampling comprised the aquatic root and the shoot during wintertime (inundation of the basin of the vernal pool), and the terrestrial root and shoot during springtime, respectively, therefore, a final number of 90 tissue samples were used. In addition to plant tissue samples per vernal pools, parallel collections of soil and water were included in the study to obtain a panoramic view of the microbial dynamics across the ecosystem layers, from soil, to water column and plant tissues.

**Table 1.** Sampling sites at Merced Vernal Pools and Grassland Reserve (MVPGR). Each site belongs to a single vernal pool, with a distance of one kilometer from each other. Three soil types are represented: Corning, Redding, and Hopeton.

| <u>Collection ID</u> | <u>Site ID</u> | <u>Distance from</u><br><u>urbanization</u> | <u>Soil parent</u><br><u>material</u> | <u>Longitude</u> | <u>Latitude</u> |
| --- | --- | --- | --- | --- | --- |
| MONT-0VP4 | K0 | 0 Kilometers | Corning | -120.42011 | 37.3788671 |
| MONT-1VP4 | K1 | 1 Kilometer | Redding | -120.41254 | 37.33804231 |
| MONT-2VP2 | K2 | 2 Kilometers | Redding | -120.40515 | 37.3878441 |
| MONT-3VP1 | K3 | 3 Kilometers | Corning | -120.38972 | 37.3898201 |
| MONT-4VP5 | K4 | 4 Kilometers | Corning | -120.37992 | 37.3889871 |
| MONT-5VP1 | K5 | 5 Kilometers | Hopeton | -120.376 | 37.3927671 |

### 1.3 Sterilization of tissues and library preparation

Sterilization was performed at the laboratory following a mechanical procedure: two rinses with distilled water for two minutes each, to separate the soil aggregates from the biological sample. The following rinses were then used: ethanol 75%, one rinse for one minute; Milli Q water, one rinse for two minutes; bleach 5%, once rinse for one minute; and Milli Q water, one rinse for five minutes (Guzman, et al 2020). After the sterilization, roots and shoots were separated with an autoclaved blade, fast-frozen by liquid nitrogen, and pulverized using a mortar. From the pulverized plant tissue, 0.2g was measured and processed with a Qiagen PowerSoil^®^ DNA extraction kit. A total of 95 extracted DNA aliquots were obtained from tissues for sequencing the 16S rRNA gene. In separate, soil samples and water samples collected to compare with plant tissue samples, were processed following the protocol in Montiel-Molina, et al. (2021), for a total of 10 extracted DNA aliquots, 5 from soil samples and 5 from water samples.

Prior to PCR amplification and library preparation, extracted DNA aliquots were analyzed for concentration and purity via Qbit^®^ technology. Peptide nucleic acid (PNA) clamps to prevent the amplification of plant mitochondria and chloroplast DNA were added to each PCR master mix (Lundberg et al, 2013). A 50ul PCR mix was prepared for each sample as follows: water 31.8 ul; Buffer 1x 31.8ul; dNTP(0.2uM) 1ul; 926R_Adapter (0.4uM) 2ul; PNA chloroplast (1.2uM) 0.6ul; PNA mitochondria (1.2uM) 0.6ul; BSA (0.2 mg/ml) 1ul; Taq(0.1 U/ul) 1ul; Barcode primers (10uM) 2ul; DNA template 5ul. Barcoded primers targeting the 16s rRNA V4-V6 coding regions (515F_915R) were used for the analysis of the bacterial endophytes (Quince, et al 2011; Apprill, et al 2015; Parada, et al 2016). The thermocycler was programmed as follows: 94°C for 3 min; 35 cycles at 94°C for 45 sec; 78°C for 10 sec; 50°C for 30 sec; 72°C for 1 min; 72°C for 10 min and then held at 4°C. Final aliquots were analyzed for DNA concentrations using a Qbit^®^ and pooled together for purification. Sequencing was achieved on the Illumina MISeq^®^ at UC Davis Genome Center. Soil and water samples were prepared and analyzed at Argonne National Lab, Chicago.

### 1.4 Bioinformatics and diversity analysis

The platform Qiime2-v.2022.2 (Bolyen et al, 2019) was used for microbiome bioinformatics analyses of plant tissues, soil and water samples, and preliminary microbiome analysis. Sequence reads were processed with DADA2 after demultiplexing, to filter chimeric reads from the analysis and to infer amplicon sequence variants (ASVs) (Callahan et al, 2016). The SILVA database was used to assign taxonomic ranks (Quast et al, 2013). Chloroplast and mitochondria sequences were filtered from all analyses. Low sequence reads were found in terrestrial samples (shoots) from the sites KM4 and KM5, and these samples were removed from further analysis.

For plant samples we assessed diversity across vernal pool compartments, roots and shoots and plant host morphology (aquatic and terrestrial, water and soils samples. Analysis was made with R studio (v.1.3.1093) using the package Qiime2R-v0.99.1 (Bisanz, 2018) to manage artifacts produced with Qiime2. Diversity and community composition analysis, comprising alpha diversity, relative abundances, distance metrics (Bray-Curtis, Unifrac), and dissimilarity analysis between samples (PCA, ADONIS-PERMANOVA) were performed using Qiime2-v2022.2 (Bolyen et al, 2019), Phyloseq-v1.36.0 (McMurdie & Holmes, 2013) and Microbiome-v1.14.0 (Lahti et al, 2017). Pearson correlations and Mantel tests were performed with Vegan 2.6-4 (Oksanen et al, 2022), to assess the effect of environmental properties, the distance between sampling sites and anthropogenic disturbance on microbial community composition. Heatmaps for abundant taxa within tissue samples were produced with Phyloseq-v1.36.0 (McMurdie & Holmes, 2013). We blasted sequences from the top 30 ASVs against the NCBI database to determine their identity and potential function.

Analysis of the core microbiome of roots were considered given data robustness in diversity analysis. Data from plant roots samples and additional soil and water samples were processed with the R package Microbiome-v1.14.0 (Lahti et al, 2017), and graphical representations were made with the Venn diagram in Euleer-v7.2.2 (Larsson, 2022). The core microbiome was defined as taxa present in least 90% of the samples. Core microbiome sequences were blasted against the NCBI database.

## 2 Results

We observed clear differences in the diversity of microbial communities across the plant life cycle (aquatic and terrestrial) and plant compartments (root and shoot). Proteobacteria and Bacteroidetes taxa dominated the community of microbial endophytes across both plant life cycle and compartments, but with multiple patterns in community dynamics (see below). Samples varied greatly in the number of ASVs present, with rarefaction curves reaching asymptotes at a sampling depth below 1000 sequences (Supplementary Fig. 1).

### 2.1 Representative taxa in amphibious plants

Samples varied substantially in taxa composition. At the phylum level, Bacteroidetes and Proteobacteria dominated the microbiome of *E. castranse*, followed by Acidobacteria, Cyanobacteria, Firmicutes, Verrucomicrobia and Tenericutes (Fig 3). The classes Alphaproteobacteria, Betaproteobacteria, Flavobacteria, Gammaproteobacteria, Saprospirae, and Sphingobacteria were the most abundant across plant compartments. Taxa from the phylum Bacteroidetes were present mostly in roots and we observed an evident shift in taxa abundance between the aquatic stage and terrestrial stage. Within this phylum, Chitinophagaceae, Cytophagaceae, Flavobacteriaceae and Sphingobacteriaceae families were represented in roots samples, and Weeksellaceae was only present in roots samples of the terrestrial morphological stage. Proteobacteria were represented by Alphaproteobacteria, Betaproteobacteria, Deltaproteobacteria, Epsilonprotreobacteria, and Gammaproteobacteria; roots harbored the most taxa in these classes in aquatic and terrestrial samples. Firmicutes and Actinobacteria were mostly present in terrestrial roots, whereas phyla such as Verrucomicrobia and Terenicutes were present in all plant samples. Firmicutes of the class Clostridia were detected mostly in samples from roots in the aquatic stage, with taxa in the family Veillonellaceae, Clostridiaceae, Bacillaceae, Peptostreptococcaceae and Lachnospiraceae.Taxa in the phylum Acidobacteria belonged to the families Bryobacteraceae, Acidobacteriaceae, Holophagacea, Koribacteraceae, Solibacteraceae, Ellin6075, MV65 and OPB3, mostly present in samples representing the aquatic stage in roots, and absent in shoots; in contrast terrestrial stage only allocated taxa in the family Acidobacteriaceae. We detected unknown taxa from the phylum Cyanobacteria mostly in shoots samples of the terrestrial stage. Taxa from the Phormidiaceae family were detected in shoots of the aquatic phase and taxa from the family Pseudanabaenaceae were detected in aquatic roots.

**Figure 3.**
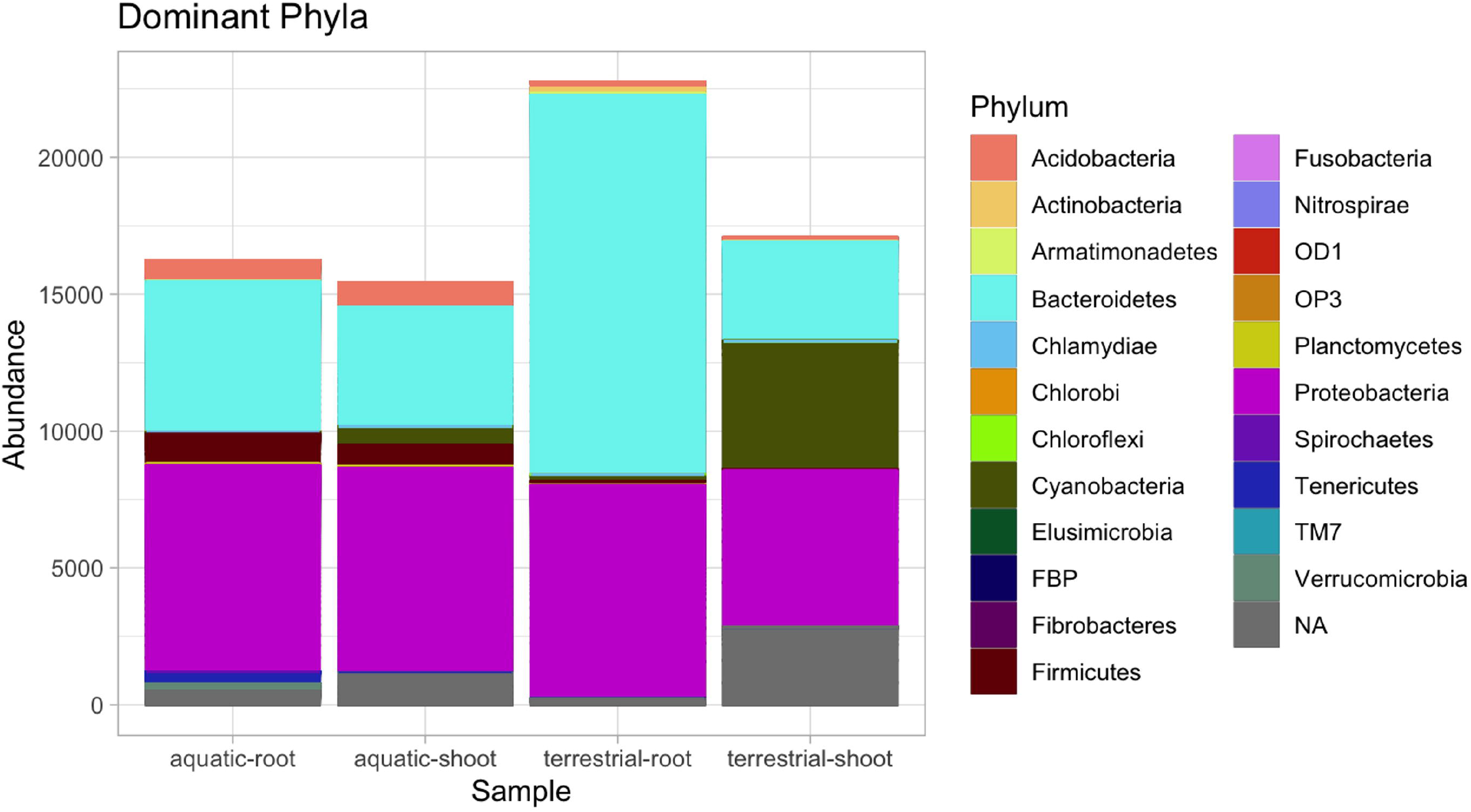
Dominant bacterial phyla in the amphibious plant E. castrense.

### 2.2 Microbial endophytic communities across plant compartments and plant morphology

Community dissimilarity among samples was analyzed using Unweighted-Unifrac and Bray-Curtis metrics, and the significance of sample clustering was corroborated by the ordination method Principal Component Analysis (Fig. 4) and PERMANOVA-ADONIS (Table 2). The samples were best defined by aquatic and terrestrial morphological stages of the plant specimens, followed by roots and shoots. The ordination analysis resulted in the formation of distinct clusters. One cluster consisted of samples corresponding to roots in the terrestrial morphological stage, while another cluster consisted of samples with roots in the aquatic morphological stage. Additionally, there was a cluster composed of samples defined by shoots in the terrestrial morphology, and a diffuse cluster formed by shoots in the aquatic morphological stage. PERMANOVA-ADONIS of Bray-Curtis and Unweighted-Unifrac dissimilarity indexes validated a significant differentiation between plant compartments and morphological stages (Table 2).

**Table 2.**
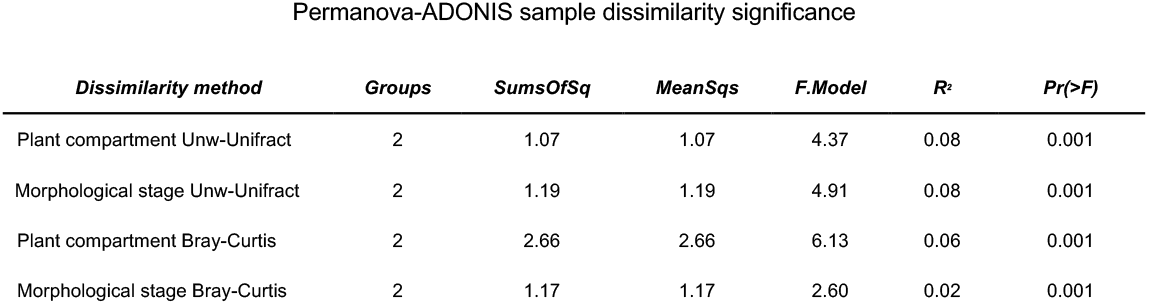
PERMANOVA-ADONIS based on microbial community composition of aquatic roots, aquatic shoots, terrestrial shoots, and terrestrial roots.

**Figure 4.**
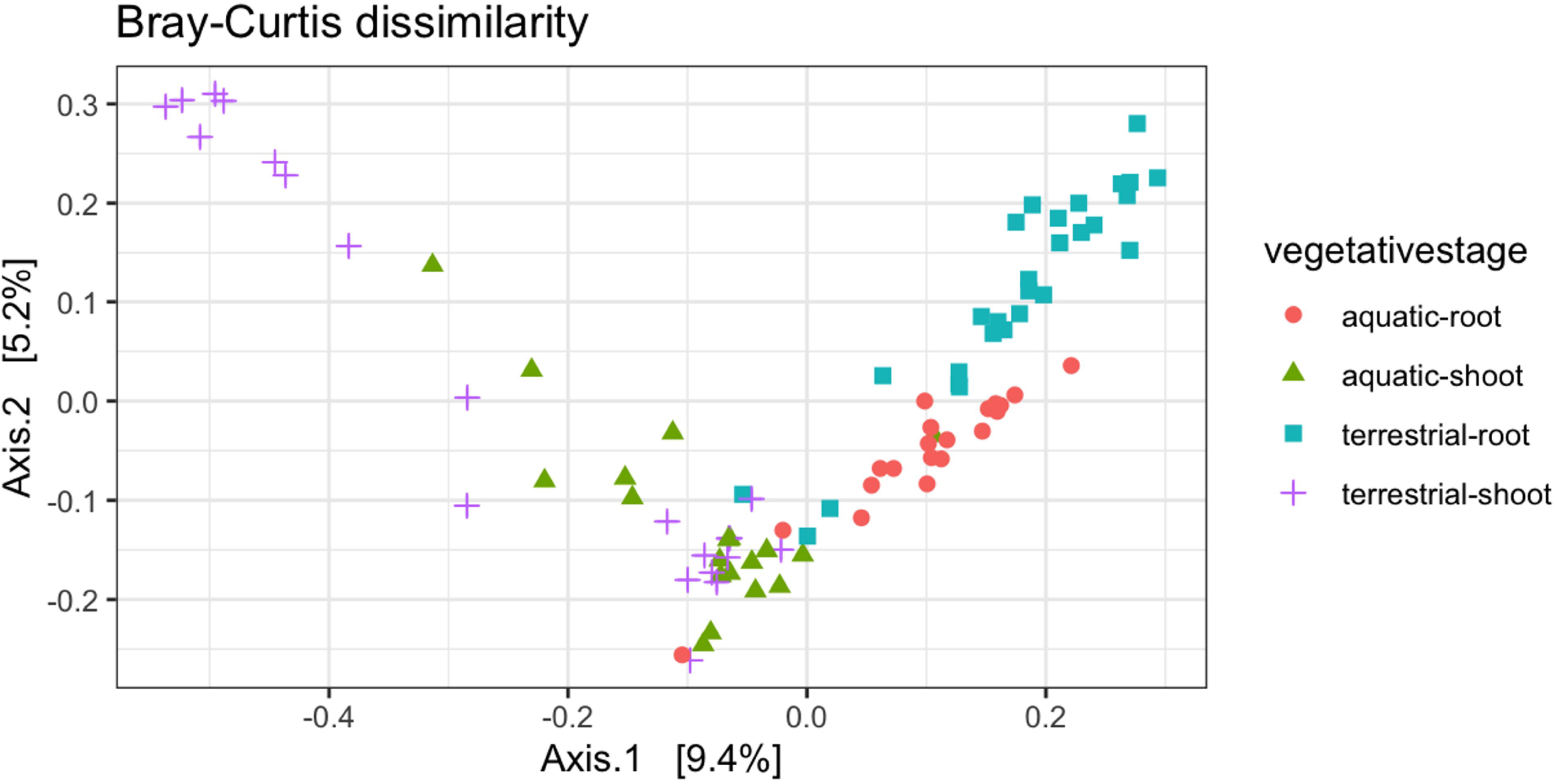
Principal Components Analysis (Bray-Curtis distances) for bacterial communities illustrates differences among plant compartments, as well as aquatic versus terrestrial phase. Samples are grouped and color shaded based on the combination of these factors.

Alpha diversity, as measured by richness, Shannon index, Simpson’s index, ACE (abundance-based coverage estimators), and Simpson’s evenness, also displayed differences between roots and shoots and between terrestrial and aquatic morphological stages. Root samples in both aquatic and terrestrial stages had higher diversity in contrast with shoots (Fig. 5). Aquatic stages showed higher overall diversity than terrestrial stages. Based on the Kruskal-Wallis test, the difference in diversity between sample types was significant (Table 3).

**Table 3.** Kruskal-Wallis non parametric analysis on alpha diversity metrics between roots and shoots samples, aquatic and terrestrial morphological stages.

| *Kruskal-Wallis Analysis* | *Shannon* | *Simpson* | *ACE* | *S evenness* | *Richness* |
| --- | --- | --- | --- | --- | --- |
| Aquatic / Terrestrial | .003 | .001 | .04 | .81 | .04 |
| Roots / Shots | 6.65<sup>-11</sup> | 1.37<sup>-11</sup> | 3.57<sup>-12</sup> | .02<sup>-3</sup> | 3.25<sup>-12</sup> |

**Figure 5.**
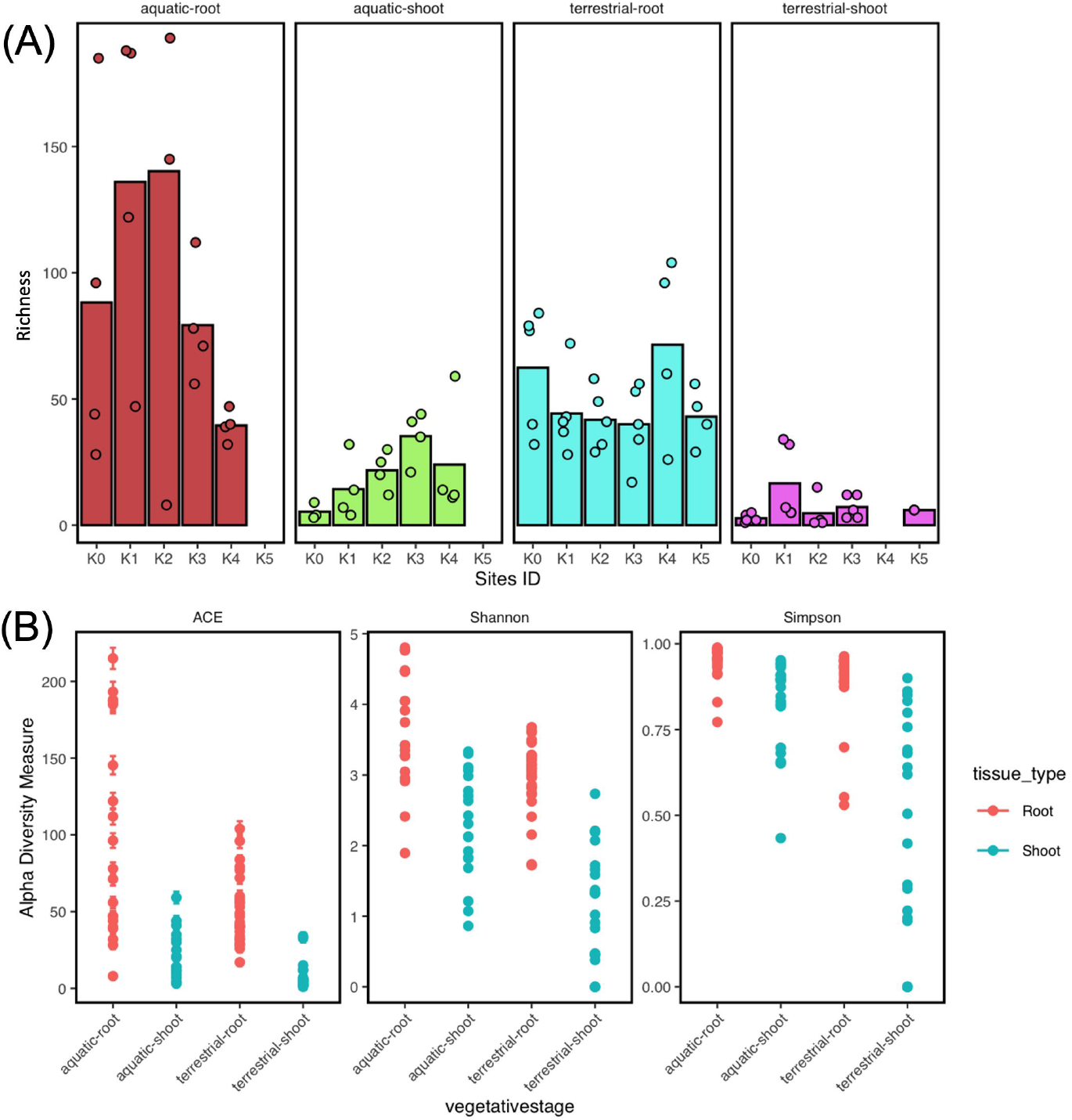
Alpha diversity in aquatic and terrestrial roots and shoots. **(A)** Differences in endophytic taxa diversity. Bars represent the total taxa abundance across sampling sites, given a specific tissue type and plant morphological stage. **(B)** Alpha diversity metrics of roots and shoots in both, aquatic and terrestrial. Dots represent individual samples.

The 30 most abundant sequences (Fig. 6) displayed varying percent sequence identity compared with taxa in the NCBI database, nevertheless in ecological terms, the composition reflected consistent niche differentiation according to plant compartment and between terrestrial and aquatic stages. For instance, Cytophagaceae spp, Chitinophaga spp, Oxalobacteraceae spp, Flavobacterium frigidarium, *Chitinophagia arvencicola, Cryseobacterum aquaticum, Pantoea agglomerans, Niastella soli* among a few other taxa within the families Mucilaginibacter, and Chitinophagia showed higher relative abundance in root tissues in both aquatic and terrestrial stages. Shoots harbored fewer of these taxa, however, terrestrial shoots had relatively high abundances of *Pantoea agglomeran*s, *Xanthomonas translucens*, and *Uzinura diaspidicola* (Fig. 6).

**Figure 6.**
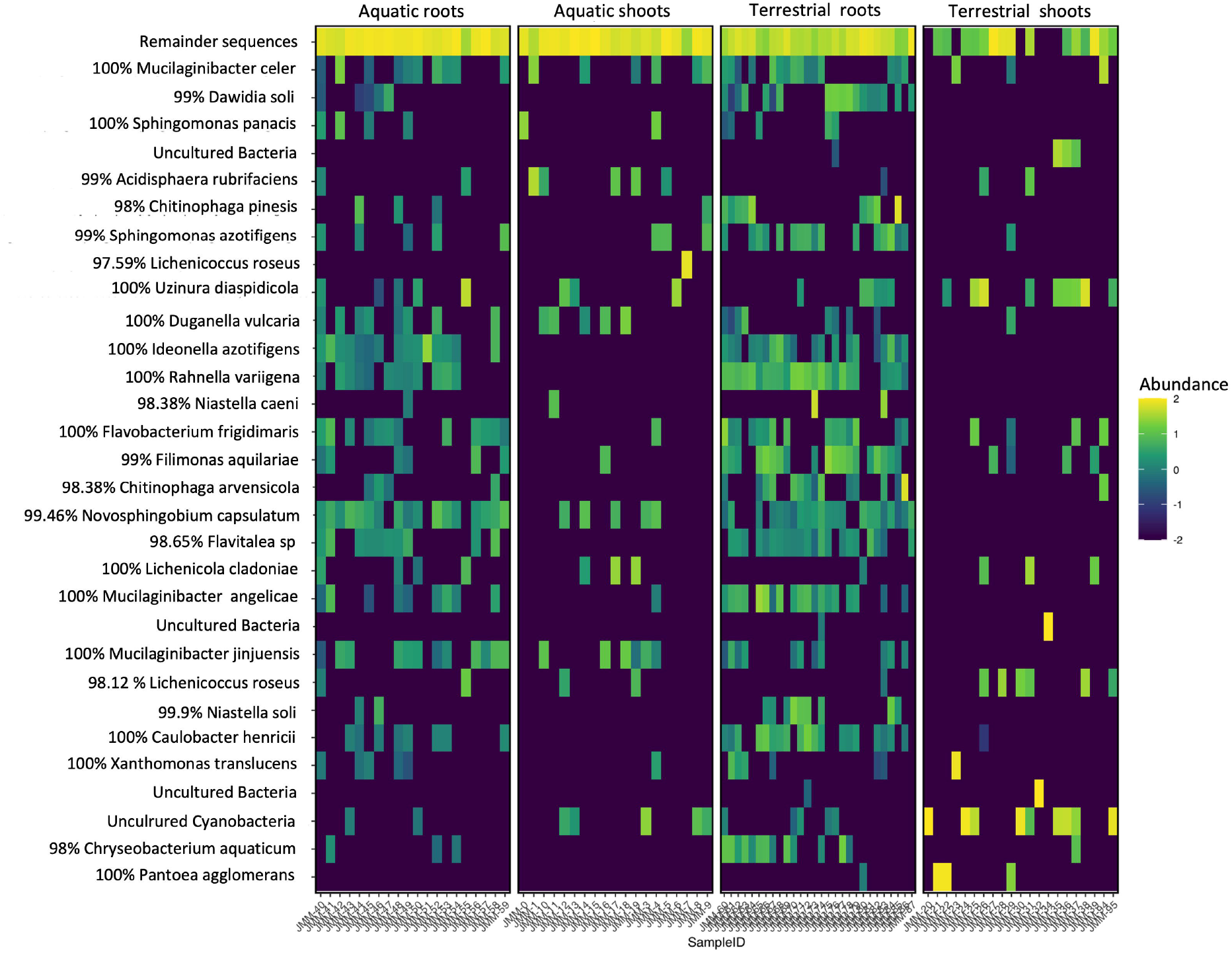
Heat map displaying the relative abundance of the 30 most abundant endophytic ASVs identified for the amphibious plant species *E*.*castrense*, organized by plant compartment. Percent identity to the closest match in NCBI is shown next to the assigned taxa name.

#### Impact of water, soil, environmental factors, and geographic distance on the plant microbiome

In contrast to the significant influence of plant compartments and plant morphology over the endophytic community, Pearson correlations showed no significant influence from the landscape spatial distances, anthropogenic activities, or soil properties. The Mantel test showed no significant distance-decay relationship between plant sample community dissimilarity and geographical distance between sites (supplementary Table 1).

Comparisons between communities in the plant endosphere and those in the surrounding water and soil environment showed differentiation. Beta diversity analysis showed habitat differentiation between samples of aquatic root morphology, terrestrial root morphology, soil and water from vernal pools ecosystems (Supplementary Fig. 2). We analyzed the core microbiome from each of these habitats; aquatic and terrestrial root tissues, water and soil (Table 4). Twenty one taxa represented the core microbiome in soil samples, and zero taxa in the case of water samples; six taxa represented the root endosphere (including aquatic and terrestrial stages), two taxa specifically belong to aquatic roots and three taxa belong to terrestrial roots, and only one taxon was shared between aquatic and terrestrial roots (Fig. 7).

**Table 4.** Core microbiomes in roots, and soil samples, and percent identity with their closest match in the NCBI database.

| Roots |  | Soil |
| --- | --- | --- |
| *terrestrial* | *aquatic* |  |
| Duganella sp. [100%] | Povalibacter sp. [95.71%] | Ktedonobacter sp. [94.39%] |
| Caulobacter rhizosphaerae [100%] | Methylovorus glucosotrophus [100%] | Psychrosinus sp. [98.13%] |
| Rhizobium cellulosilyticum [100%] | Novosphingobium sp. [100%] | Candidus Koribacter sp. [97.33%] |
| Novosphingobium sp. [100%] |  | Uncultured Acidobacteria 1 |
|  |  | Uncultured Acidobacteria 2 |
|  |  | Koribacter versatilis [98.76%] |
|  |  | Uncultured Actinomycete 1 |
|  |  | Uncultured Actinomycete 2 |
|  |  | Uncultured Acidobacteria 3 |
|  |  | Ramlibacter ginsenosidimutans [100%] |
|  |  | Acidovorax avenae [100%] |
|  |  | Uncultured bacterium 1 |
|  |  | Psicinibacter sp. [100%] |
|  |  | Bradyrhizobium japonicum [100%] |
|  |  | Anaeromyxobacter sp.[98.7%] |
|  |  | Anaeromyxobacter diazotrophicus [99.1%] |
|  |  | Uncultured bacterium 2 |
|  |  | Geomonas subterranea [100%] |
|  |  | Uncultured Deltaproteobacteria [99.8%] |
|  |  | Uncultured Acidobacteracea [98.38%] |
|  |  | Uncultured Actinobacteria [98.39%] |

**Figure 7.**
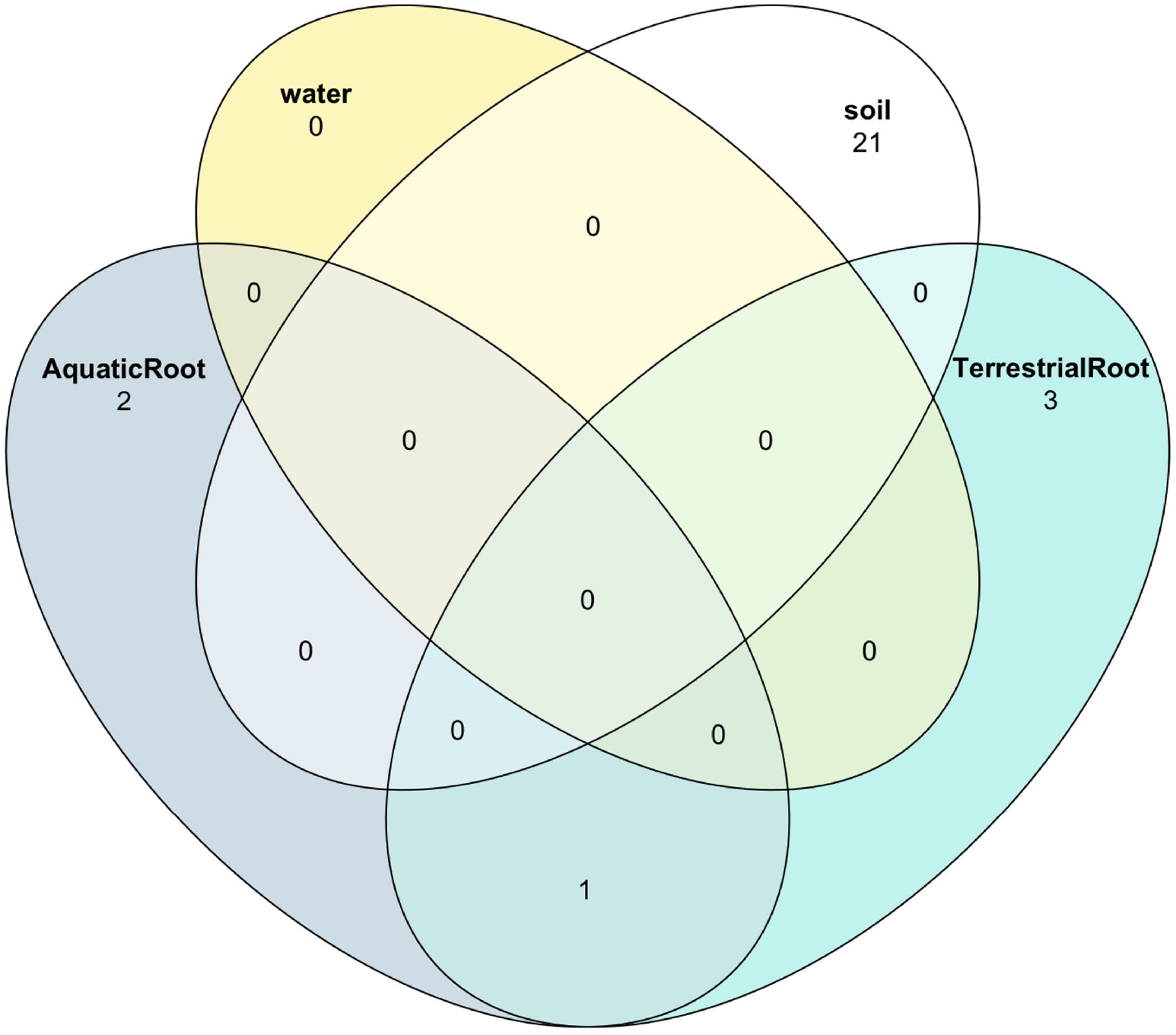
Overlap in core microbiome by ecosystem compartment in vernal pools.

## 3 Discussion

Plant species with an amphibious lifestyle are an interesting subject of study given their adaptations to contrasting conditions. These amphibious plants are mostly restricted to ephemeral wetlands known as vernal pools in Mediterranean climate regimes (Keeley and Zedler, 1998), where they have adapted to water deficit, which for some species has led to differentiated morphologies throughout their life cycle (Keeley,1999). For this study, we aimed to obtain novel information from a plant-microbe symbiosis perspective by providing the first description of the microbial endophytes in amphibious plants and addressing the transitions during their life cycle. Information on microbial community assembly may be key to understanding the mechanism behind the amphibious behavior, as microbial symbionts could be involved in alleviating plant stress under desiccation conditions, with important implications for both agricultural and natural ecosystems where water becomes a limiting factor.

When vernal pool plants are young, they resemble an aquatic grass, and as they become mature their morphology is more similar to a spiny weed. We studied the dynamics of the endophytic community across the plant life cycle as a first step towards elucidating the potential role of microbes in alleviating the stress triggered by the inundation and conditions experienced by the plants. The results obtained from the beta diversity metrics defined two well-delimited niches according to aquatic and terrestrial stages and the morphology of the amphibious plants. Our result supported the hypothesis that the endophytic community varies by plant stage, and across the life cycle of the plant. Our results are consistent with Zhou et al (2023), who reported significant differences in the microbiome of young and mature specimens of *Suaeda salsa*, a plant inhabiting high salt concentration habitats. Our study yielded contrasting results regarding water stress compared to other studies—but this is likely due to the more pronounced differences in water availability captured in our study compared to the more limited differences examined previously. For example, in Eucalyptus trees, water deficit treatments were not a significant driver of microbial community variation (Dasgupta et al, 2020), and a study of pepper plants roots and fruits microbiomes reported that bacterial endophytes were not affected by water irrigation limitation (Cui, et al 2020). Both studies suggested that endophytic communities were resilient to water stress. Yet, in our study, plant stage (aquatic vs terrestrial) was a stronger driver of microbial community variation than compartment, supporting our hypothesis that endophytic communities of vernal pool plants shift significantly with the extremes that plants experience in vernal pool ecosystems. This may suggest a role for bacterial endophytes in alleviating part of the stress generated by the water deficit and inundation conditions within vernal pools. This result also suggests that amphibious plants have the capacity to select their symbionts. However, inoculation experiments will be required to test if recruitment of symbiotic microbial communities is part of vernal pool plant adaptation to the contrasting conditions of inundation and desiccation.

Differences in microbial communities between compartments is generally attributed to the presence of differential bioactive compounds. For example, Toju et al. (2019) suggested that in tomato plants, alkamides, caffeic acid derivatives, polysaccharides and alkenes affect the microbiome. We did not analyze secondary metabolites in our study, but the taxa associated with above- and below-ground provide an approach to understand ecological relations (Table 5). Other reasons can include unhealthy conditions (with dark green leaves, signs of grazing) that diminish the diversity in the tissues (Yang et al, 2022). Finally, the niche differentiation can be also attributed by the lack of free entry of endophytes into living tissues in roots and shoots, which might require bacterial capabilities to hydrolyse the hydrophobic incrustations of the walls of epidermal, hypodermal, endodermal and other cortical cells (McCulley, 2001).

**Table 5.** Ecological role of top ASVs associated with amphibious plants.

| Bacterial taxa | Ecological role | Niche | % NCBI Identity | References |
| --- | --- | --- | --- | --- |
| *Mucilaginibacter celer* | Improves growth and salt tolerance, some isolates also from freshwater ecosystems | All | 100% | Kim et al, 2020 |
| *Dawida soli* | Heterotrophic aerobic metabolism, endophytes of roots | Roots terrestrial and aquatic | 99% | Manter et al, 2010 |
| *Sphingomonas panacis* | Heavy metals resistance, rhizosphere resident | Aquatic roots, shoots and terrestrial roots | 100% | Kim et al, 2019 |
| *Acidisphaera rubrifaciens* | Acidic environments, aerobic, primary producer [bacteriochlorophyll-containing] | Aquatic roots, shoots and terrestrial shoots | 97.86% | Hiraishi et al, 2000 |
| *Chitinophaga pinesis* | Chitin degradation, fungal cell walls degradation | Roots | 98% | Lu et al, 2023 |
| *Sphingomona azotifigens* | Nitrogen fixing bacteria, growth promoter | Aquatic roots, shoots and terrestrial roots | 99% | Videira et al, 2009 |
| *Lichenicoccus roseus* | Antarctic lichen symbiont, xylan decomposer | Aquatic shoots | 97.59% | Pankratov et al, 2020 |
| *Uzinura diaspidicola* | Insects symbiont, nutrient acquisition | All | 100% | Sabree et al, 2012 |
| *Duganella vulcaria* | Endophytes roots and shoots, growth promoter | Aquatic roots, shoots and terrestrial roots | 100% | Baldani et al, 2014 |
| *Ideonella azotifigens* | Rhizosphere, nitrogen fixer | Roots terrestrial and aquatic | 100% | Noar & Buckley, 2009 |
| *Rahnella variigena* | Endophyte, growth promoter | Roots terrestrial and aquatic | 100% | Raj et al, 2023 |
| *Filimonas aquilariae* | Plant roots resident, high CO<sub>2</sub> dependent | All | 99% | Lin et al, 2017 |
| *Chitinophaga arvensicola* | Wetlands, plant detritivorous | Roots terrestrial and aquatic | 98.38% | Pankratov et al, 2006 |
| *Novosphingobium capsulatum* | Rhizosphere inhabitant, Sulfuric metabolism: alkaline sulfonate assimilation | Aquatic roots, shoots and aquatic shoots | 99.46% | Kumar et al 2017 |
| *Flavitalea* sp. | Root endophyte | Roots terrestrial and aquatic | 98.65% | Fernandez-Gonzalez et al 2019 |
| *Lichenicola cladoniae* | Cold adapted lichen symbiont (Antarctica) | All | 100% | Noh et al, 2020 |
| *Mucilaginibacter angelicae* | Rhizosphere, cellulose-degrading | Roots terrestrial and aquatic | 100% | Kim et al 2012 |
| *Mucilaginobacter jinjuensis* | Rotten wood xylan-degrading, latent in plant roots | Roots terrestrial and aquatic | 100% | Khan et al, 2016 |
| Niastella soli | Rhizosphere, denitrifier | Roots terrestrial and aquatic | 99.9% | Akter et al, 2021 |
| Caulobacter henricii | Fresh water and rhizosphere, associated with caulophages "Kronos" | Roots terrestrial and aquatic | 100% | Berrios and Ely, 2019 |
| Xanthomonas translucens | Plant pathogen | Roots terrestrial and aquatic | 100% | Sapkota et al, 2020 |
| Flavobacterium frigidarium | Sediments in Antarctica, detritivorous of plants and algae | Roots and less proportion shoots | 100% | Humphry et al, 2001 |
| Chryseobacterium aquaticum | Fresh water, see also Elizabethkingia | Roots terrestrial and aquatic | 98% | Bernardet et al, 2006 |
| Pantoea agglomerans | Associated with plant roots. It could be associated with human infections via wound infection with plant material. | Terrestrial shoots | 100% | Cruz et al, 2007 |

The microbial endophytic communities in our study were dominated by Bacteroidetes and Proteobacteria (Fig. 3), comparable to the endophytes in Eucalyptus, which require large amounts of water (Dasgupta et al, 2020) and aquatic habitats defined by low tides and high oxygen availability (Stevens et al, 2005). The Proteobacteria phylum is ubiquitous in both aquatic and terrestrial ecosystems, and endophytic taxa from this phylum fill important niches in relation with biogeochemical cycles such as nitrogen (Zhou et al, 2020, Bai et al 2022). Bacteroidetes are likewise involved in a variety of processes, including pathogen suppression (Lidbury et al, 2020). In our study, we observed taxa related with phytopathogens (although no symptoms of disease were noticed at the moment of handling specimens) and nitrogen fixers in roots during the aquatic stage of the plants, similar to rice crops (Verma et al, 2001). We summarize in Table 5 information about the top ASVs of endophytes to provide insight into their potential ecological role in amphibious plants.

Beyond niche differentiation by plant compartment, we examined the surrounding soil and water to compare microbiomes between these ecosystemic layers. These ecosystem compartments (soil, water and plant tissues) showed strong dissimilarities in bacterial community composition. Other studies have reported that endosphere, rhizosphere and soil hold a niche differentiation, and a transition from soil, rhizosphere, rhizoplane and plant endosphere, where diversity is higher outside the plant tissues (Wang et al, 2022). In our study, soil samples exhibited higher microbial diversity in comparison with plant tissues (supplementary Figure 3), and we detected a similar pattern when tissues were compared with water samples, however these samples were more variable than soil.

## 4 Conclusion

In Mediterranean ecosystems where summer is characterized by desiccation and winter by inundation, amphibious plants specially adapted to conditions of stress are hypothesized to have microbial symbionts with the potential to alleviate water stress. This study shows that in the amphibious vernal pool plant *E. castranse*, plant host morphology and tissue compartmentalization has strong influence over the bacterial endophytic community assembly. Future research on microbial endophytes in amphibious plants should incorporate the assessment of the fungal endophytes as well, as fungi play a significant role in the plant microbiome and have unique ecological functions.

This pioneering study on the microbiome of vernal pool amphibious plants provides preliminary findings about the role of endophytes in ecosystems subject to environmental change events, with parallels in other systems in our changing world. The interactions and mechanics underlying this microbial-plant association is complex, with endophytic communities also occurring during periods of variation in oxygen concentrations and other effects within the environment not examined here. Future research should incorporate those other conditions, which have effects in other ecosystems and with other organisms. Presently, there is a growing interest in artificial environments that are resilient and have a circular energy flow. The microbiome of amphibious plants can be an interesting addition to research focusing on such technology. We propose that future research on the benefit of endophytes to amphibious plants involve an experimental approach, where plants are inoculated with endophytes and tested for survival in growing chambers, and we suggest the use of specimens within the same family plants that are used as crops as we did here, using *E. castrense*, a member of the carrot family. The importance of these following steps will be useful for the implementation of agricultural systems.

We suggest exploring plant relatives from the carrot family for further studies, as the data obtained in this study can serve as a foundation for future research involving crop plants from the same family.

## 7 Acknowledgments

We knowledge the participation of Dr. Iolanda Ramalho da Silva, Andrea Salinas, David Pérez, Kehily Maldonado, Kimberly Du and other members from Frank, Sexton and Beman’s labs. CONACyT-UCMexus, institutions who provided with financial support for Dr Montiel-Molina during his tenure as a doctoral student. To the California Native Plant Society who provided financial support, and UC Merced Natural Reserve System allowing to perform the research at the UCMerced Vernal Pools and Grassland Reserve.

## Supplementary Material

### 1 Supplementary Tables

**Supplementary Table 1.** Mantel test of spatial distance (geo position) and diversity metrics.

| *Spatial distance Mantel* | <u>Observed</u> | <u>Shannon</u> | <u>Simpson</u> | <u>ACE</u> |
| --- | --- | --- | --- | --- |
| Statistic r | -0.04 | -0.04 | 0.06 | -0.04 |
| Significance | 0.8 | 0.8 | 0.08 | 0.8 |

**Supplementary Table 2.** PERMANOVA-ADONIS by habitat type in vernal pool ecosystems, Root endosphere versus soil and water.

Permanova-ADONIS sample dissimilarity significance
| *Dissimilarity method* | *Groups* | *SumsOfSq* | *MeanSqs* | *F.Model* | *R<sup>2</sup>* | *Pr(>F)* |
| --- | --- | --- | --- | --- | --- | --- |
| Unw-Unifract | 3 | 4.05 | 1.35 | 5.77 | 0.24 | 0.001 |
| Bray-Curtis | 3 | 3.8 | 1.26 | 3.38 | 0.16 | 0.001 |

### 2 Supplementary Figures

**Supplementary Figure 1.**
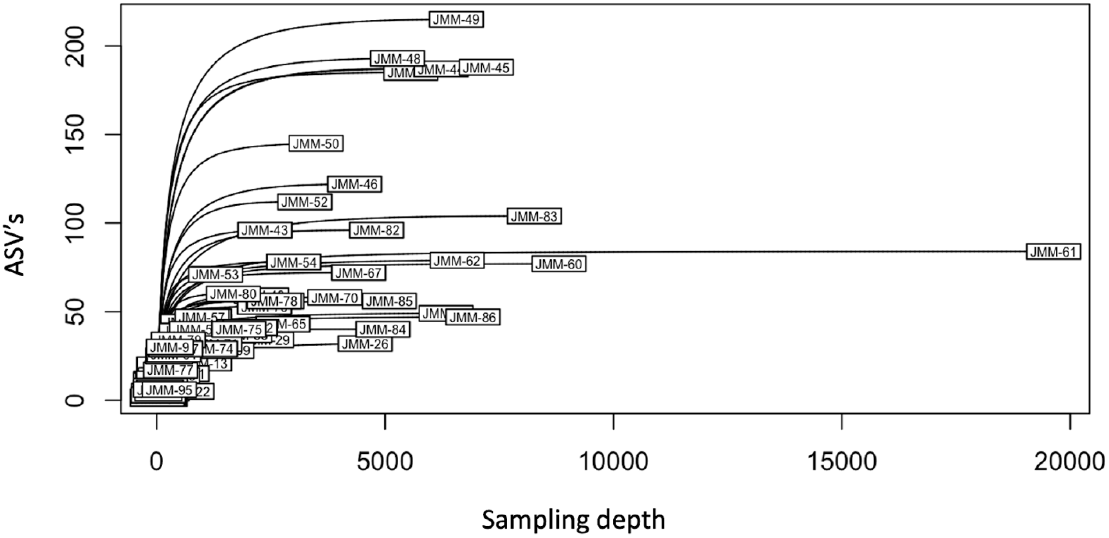
Sequence variants (ASV’s) identified by the number of ASVs.

**Supplementary Figure 2.**
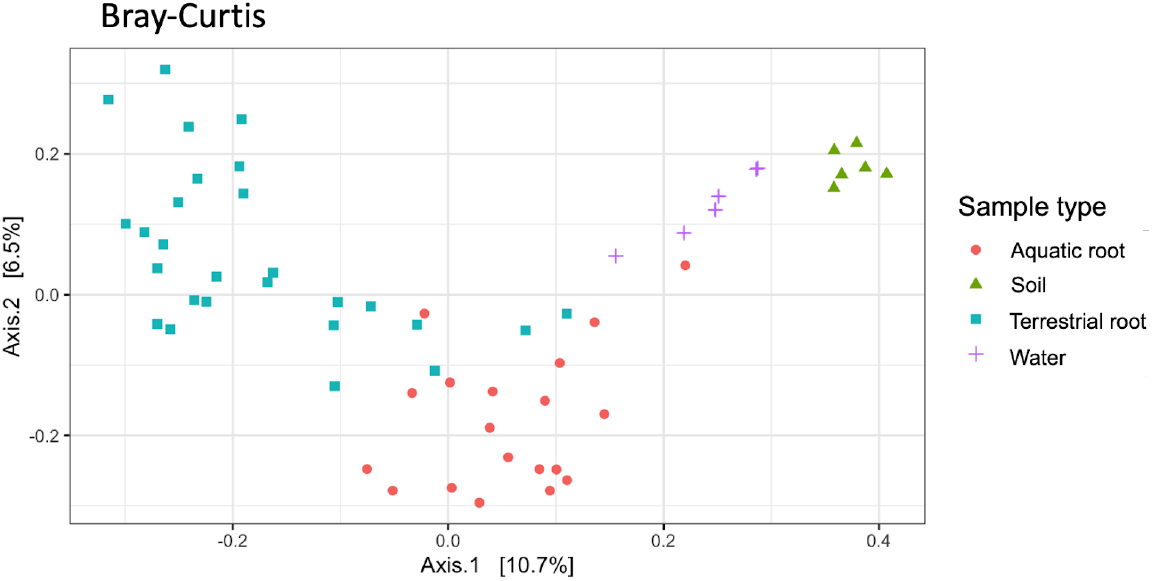
PCA based on Bray-Curtis dissimilarity index between tissue samples, water and soil samples.

**Supplementary Figure 3.**
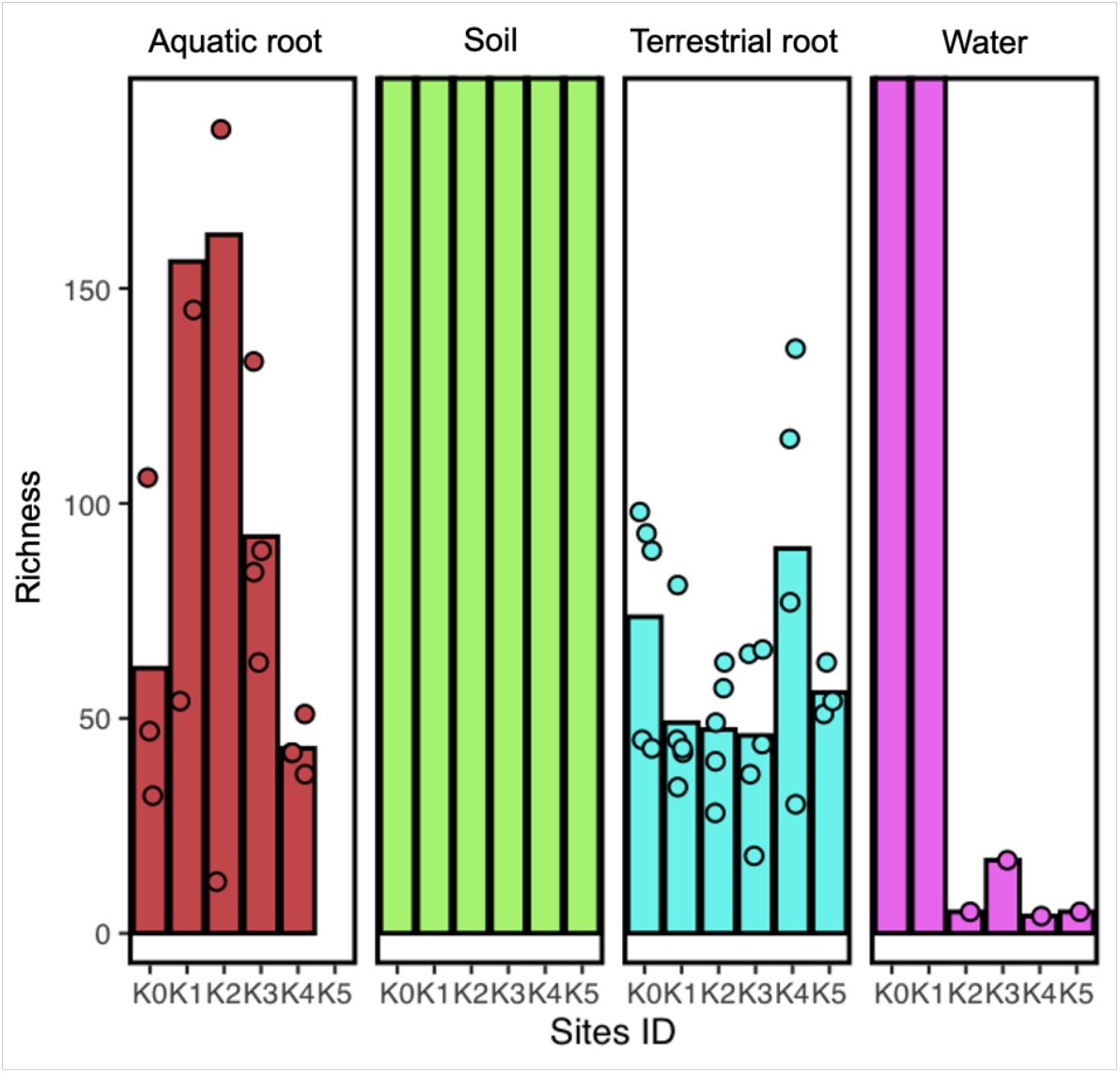
Differences in diversity between ecosystem compartments, soil, water, and plant tissues.

